# Ceftriaxone and mGlu2/3 interactions in the nucleus accumbens core affect the reinstatement of cocaine-seeking in male and female rats

**DOI:** 10.1101/2020.02.17.952762

**Authors:** Carly N. Logan, Allison R. Bechard, Peter U. Hamor, Lizhen Wu, Marek Schwendt, Lori A. Knackstedt

**Author notes:** Corresponding author: Lori A. Knackstedt, Department of Psychology, University of Florida, 945 Center Dr., Gainesville, FL, 32611.

## Abstract

**Rationale:** The beta-lactam antibiotic ceftriaxone reliably attenuates the reinstatement of cocaine-seeking. While the restoration of nucleus accumbens core (NA core) GLT-1 expression is necessary for ceftriaxone to attenuate reinstatement, AAV-mediated GLT-1 overexpression is not sufficient to attenuate reinstatement and does not prevent glutamate efflux during reinstatement.

**Aims:** Here, we test the hypothesis that ceftriaxone attenuates reinstatement through interactions with glutamate autoreceptors mGlu2 and mGlu3 in the NA core.

**Methods:** Male and female rats self-administered cocaine for 12 days followed by 2-3 weeks of extinction training. During the last 6-10 days of extinction, rats received ceftriaxone (200 mg/kg IP) or vehicle. In experiment 1, rats were killed, and NA core tissue was biotinylated for assessment of total and surface expression of mGlu2 and mGlu3 via western blotting. In experiment 2, we tested the hypothesis that mGlu2/3 signaling is necessary for ceftriaxone to attenuate cue- and cocaine-primed reinstatement by administering bilateral intra-NA core infusion of mGlu2/3 antagonist LY341495 or vehicle immediately prior to reinstatement testing.

**Results:** mGlu2 expression was reduced by cocaine and restored by ceftriaxone. There were no effects of cocaine or ceftriaxone on mGlu3 expression. We observed no effects of estrus on expression of either protein. The antagonism of mGlu2/3 in the NA core during both cue- and cocaine-primed reinstatement tests prevented ceftriaxone from attenuating reinstatement.

**Conclusions:** These results indicate that ceftriaxone’s effects depend on mGlu2/3 function and possibly mGlu2 receptor expression. Future work will test this hypothesis by manipulating mGlu2 expression in pathways that project to the NA core.

## Introduction

Recent estimates suggest approximately 913,000 Americans meet the Diagnostic and Statistical Manual of Mental Disorder’s criteria for cocaine dependence or abuse (National Survey on Drug Use and Health, 2016). Although cocaine use disorder is prevalent, there are currently no FDA-approved treatments for this disorder. Pharmacological options to reduce relapse are necessary as even after extended periods of abstinence, there are high rates of relapse to cocaine use (O’Brien, 2003).

Animal models of drug seeking can be used to identify drug-induced neuroadaptations and potential therapies for relapse prevention. The intravenous self-administration extinction-reinstatement paradigm has been used to model cocaine relapse in rodents (Epstein et al 2006). In this model, operant cocaine self-administration is paired with cues such as a light and tone. Once self-administration criterion is met, extinction training during is conducted, during which time the conditioned behavior no longer yields drug delivery or drug-associated cues. The drug seeking response can then be reinstated via noncontingent drug delivery or presentation of the cues previously associated with drug delivery. Utilization of this model has identified the dysregulation of glutamatergic neurotransmission following cocaine self-administration as a mediator of reinstatement. Namely, there is an increase in synaptically released glutamate in the nucleus accumbens core (NA core) during cocaine- (McFarland et al 2003; Trantham-Davidson et al 2012) and cue-primed reinstatement of cocaine-seeking (Smith et al 2017).

Several alterations in the NA core contribute to the increase in synaptically released glutamate which drives the reinstatement of cocaine-seeking. Basal levels of non-synaptic glutamate are decreased in the NA core 2-3 weeks after discontinuation of cocaine self-administration (Baker et al 2002). Basal glutamate levels are maintained by system x_c_-, an antiporter that exchanges intracellular glutamate for extracellular cystine (Mcbean and Flynn 2001). Expression and function of xCT, the catalytic subunit of system x_c_-, is decreased following discontinuation of cocaine self-administration (Baker et al 2003; Knackstedt et al 2010). Expression of GLT-1, the primary glutamate transporter (Haugeto et al 1996), is also decreased in the NA core following 3 weeks of extinction from cocaine self-administration (Knackstedt et al 2010).

The β-lactam antibiotic ceftriaxone and the cystine prodrug N-acetylcysteine have been proposed to be potential treatments for cocaine relapse prevention (Baker et al., 2003; Knackstedt et al., 2010). Both compounds restore non-vesicular basal glutamate levels and prevent the increase in synaptic glutamate in the NA core during the reinstatement of cocaine-seeking (Baker et al 2003; Trantham-Davidson et al 2012). Restoration of GLT-1 and xCT expression in the NA core is necessary for ceftriaxone to attenuate reinstatement of cocaine-seeking (LaCrosse et al 2017) and restoration of GLT-1 is necessary for N-acetylcysteine to prevent reinstatement (Reissner et al 2015). However, AAV-mediated overexpression of GLT-1 in the NA core is not sufficient to attenuate reinstatement of cocaine-seeking (Logan et al 2018). Although such overexpression of GLT-1 attenuates glutamate efflux during cocaine-primed reinstatement, it does not fully prevent efflux as ceftriaxone does (Trantham-Davidson et al., 2012). Therefore, other mechanisms must account for the ability of ceftriaxone to attenuate glutamate efflux and reduce reinstatement of cocaine-seeking. Such alterations may include normalization of basal glutamate tone in the NA core and/or restored expression of receptors that regulate glutamate release, such as the G_i_-coupled group II metabotropic glutamate receptors 2 (mGlu2) and 3 (mGlu3).

Stimulation of presynaptic mGlu2 and mGlu3 receptors negatively modulates neurotransmitter release by inhibiting cyclic AMP (cAMP)-dependent phosphorylation of N-Type Ca2+ channels (Swartz and Bean 1992; Tanabe et al 1993; Conn and Pin 1997; Anwyl 1999). mGlu2/3 also regulate non-vesicular release of glutamate via system x_c_- (Baker et al 2002; Moran et al 2005). Due to their ability to regulate glutamate levels, mGlu2/3 receptors have been implicated in relapse. Systemic administration of a mGlu2/3 agonist attenuates cocaine- and cue-primed reinstatement, but also reduces food seeking (Baptista et al 2004; Peters and Kalivas 2006; Justinova et al 2016). Assessment of mGlu2 and mGlu3 protein expression after cocaine exposure was not possible prior to recent development of antibodies specific to each receptor that are used for the present studies. A decrease in mGlu2/3 receptor density in the prefrontal cortex and NA core following cocaine self-administration and extinction training has been detected using radioligand binding (Pomierny-Chamiolo et al 2017). Following cocaine self-administration there is also a decrease in NA core mGlu2/3 function, as assessed by field potential recordings in the presence of an mGlu2/3 agonist (Moussawi et al 2011).

Antagonism of mGlu2/3 prevents N-acetylcysteine from attenuating reinstatement of cocaine-seeking (Moran et al 2005; Moussawi et al 2011). However, the contribution of mGlu2/3 receptor expression and function in ceftriaxone’s ability to reduce reinstatement of cocaine-seeking has not been investigated. Here, we tested the hypothesis that cocaine self-administration followed by 2 weeks of extinction training would produce a decrease in surface and total expression of mGlu2 and mGlu3 in the NA core. Additionally, we hypothesized that in cocaine-experienced rats, ceftriaxone would restore mGlu2 and mGlu3 surface and total expression in the NA core in male and female rats. We previously found that ceftriaxone is unable to attenuate the reinstatement of cocaine-seeking in female rats tested in the estrus phase of the estrous cycle but does attenuate reinstatement when rats are in non-estrus phases (Bechard et al., 2018). Thus, here we sought to determine whether mGlu2 or 3 expression accounts for this effect. We also tested the hypothesis that activation of mGlu2/3 in the NA core is necessary for ceftriaxone to attenuate the reinstatement of cocaine-seeking in male and female rats.

## Methods and materials

### Animals

Adult male and female Sprague-Dawley rats, (300-350 and 225-275 g at surgery, respectively, Charles River, Laboratories, Raleigh, NC, USA) were housed in a vivarium with controlled temperature and humidity on a 12 h reversed-light dark cycle. All procedures were carried out during the dark phase of the cycle. Rats underwent 1 week of habituation with ad libitum access to food and water before receiving surgery. Experiment 1 included 22 male rats: 15 male rats that underwent cocaine self-administration, and 7 rats that received sham surgery and remained cocaine naive. Experiment 2 included 40 rats (27 females; 13 male) that all self-administered cocaine. After surgeries, animals were food restricted to 20 g of standard chow and permitted ad libitum access to water. All procedures were approved by the Institutional Animal Care and Use Committee of the University of Florida.

### Drugs

Cocaine-HCl (NIDA Controlled Substances Program, Research Triangle Institute, NC, USA) was dissolved in sterile 0.9% physiological saline. The concentration of cocaine was 4 mg/mL for males (0.177 mg/infusion) and 3.2 mg/mL for females (0.141 mg/infusion). Ceftriaxone was dissolved in vehicle (0.9% physiological saline) at a concentration of 200 mg/mL. For both experiments, ceftriaxone (200 mg/kg) or vehicle (1 mL/kg) was administered intraperitoneally (IP) immediately following extinction sessions on the final 6-10 days of extinction training. The range of ceftriaxone treatment length was to ensure that extinction criteria was met and that female rats were in the correct phase of the estrous cycle for sacrifice/testing. This dose of ceftriaxone has been reliably shown to attenuate cue- and cocaine-primed reinstatement in males and females (Bechard et al., 2018; Knackstedt et al., 2010; Fisher et al., 2012; Sari et al., 2009). For Experiment 2, the mGlu2/3 antagonist LY341495 (10 μM, 0.5 μL, 3.53 ng/μL) or its vehicle (0.9% physiological saline, 0.5 μL) was infused bilaterally 5 minutes before each reinstatement test.

### Surgery

For both experiments, rats were anesthetized with ketamine (males 87.5 mg/kg, IP; females 60 mg/kg, IP) and xylazine (males and females 5 mg/kg, IP) prior to surgery. Carprofen (1 mg/kg, SC) was administered as an analgesic for three days following surgery. Catheters (SILASTIC silicon tubing, ID 0.51 mm, OD 0.94 mm, Dow Corning, Midland, MI) were implanted in the jugular vein and secured via suture thread. Catheters passed subcutaneously through the shoulder region and exited the back of the animal. The catheter was secured to a cannula (Plastics One, Roanoke, VA, USA) embedded in a rubber harness (Instech, Plymouth, Meeting, PA, USA). This harness was worn for the duration of the self-administration portion of the study. Catheters were flushed daily with 0.2 mL of heparinized saline (100 IU/mL) and catheter patency was verified periodically via intravenous administration of methohexital sodium (10 mg/mL, IV), which produces a temporary loss of muscle tone.

Rats used for Experiment 2 also underwent stereotaxic implantation of bilateral guide cannulas (Plastics One, Roanoke, VA, USA) 2 mm above the NA core (AP 1.2 mm, ML 2.5 mm, DV −5.5 mm). Cannulas were secured using dental cement and stainless-steel skull screws, and a stylet was inserted into the cannula until microinjections were administered prior to the reinstatement tests.

### Estrous cycle tracking

During the last week of extinction training, the estrous cycle was monitored as previously described (e.g., Bechard et al. 2018; Marcondes et al., 2002; Lebron-Milad and Milad 2012; McLean et al., 2012). The vaginal opening was gently lavaged with physiological saline (0.9%). The saline wash was collected and placed on a slide for the immediate categorization of estrous cycle phase by visual observation of the proportion of leukocytes, nucleated epithelial cells, and cornified epithelial cells using a light microscope (10× magnification). Proestrus was characterized by a majority of nucleated epithelial cells, a majority of cornified epithelial cells indicated estrus; diestrus was characterized by a majority of leukocytes; and metestrus by an equal proportion of each of these cell types. Phases were later confirmed following the staining of samples with cresyl violet. Vaginal lavage samples were collected and classified 10 minutes prior to euthanasia/reinstatement testing.

### Cocaine self-administration, extinction and reinstatement

Cocaine self-administration sessions occurred daily between the hours of 9 am and 12 pm. For these sessions, rats were placed in operant conditioning chambers (30 × 24 × 30 cm; Med Associates, St. Albans, VT, USA), which were equipped with two retractable levers. Presses on the inactive lever were recorded but had no programmed consequences. Presses on the active lever resulted in an infusion of cocaine concurrently with a tone (2900 Hz) and illumination of a light above the active lever. Infusions were delivered on a FR1 reinforcement schedule with a 20 second time out period after cocaine delivery. During the time out period, lever presses were recorded but did not result in a delivery of cocaine or presentation of the light or tone. Presses on the active lever resulted in cocaine delivery (0.5 mg/kg/infusion in 0.05 μL). Self-administration sessions occurred daily (2 hr/day) until rats met criterion of 9 or more infusions for 12 days. After meeting criteria, rats underwent extinction training in the same operant chamber. During extinction training, presses on both levers were recorded, but had no programed consequences. Extinction sessions occurred daily (2 hr/day) for 2-3 weeks until presses on the previously active lever were <15 for 2 consecutive sessions. During the last week of extinction training, rats were treated with ceftriaxone or vehicle immediately following the extinction session for at least 6 days prior to being killed for Experiment 1 or prior to the first reinstatement test of Experiment 2. Rats in Experiment 2 underwent reinstatement tests as described below.

### Biotinylation protocol

Rats were killed 18-21 days after the cessation of cocaine self-administration and at least 6 days of Cef/Veh treatment. Rats were rapidly decapitated without anesthesia and the NA core was dissected on ice. The tissue was cut into 200 μM prism-shaped sections with a McIllewan tissue chopper (Ted Pella) and incubated in aCSF containing Sulfo-NHS-SS-Biotin (Pierce) at 4°C. The reaction was quenched with glycine buffer and sections were homogenized. A portion (250 μg) of each sample lysate was incubated overnight with streptavidin agarose beads (Sigma) to capture biotinylated proteins. The remainder of the sample was stored as the total protein fraction. On the following day, streptavidin-coated beads with attached biotinylated proteins were separated by centrifugation from the non-biotinylated proteins. Biotinylated proteins were eluted from beads with Laemmli sample buffer containing a reducing agent. Samples were stored frozen until use. The amount of mGlu2 and 3 protein in the total and surface (biotinylated) fractions were analyzed by Western blotting.

### Western blotting

Proteins were separated using 10% SDS-PAGE and transferred to PVDF membrane. The membranes were probed overnight at 4°C with primary antibodies diluted in 5% milk. Antibodies that distinctly recognize mGlu2 (Millipore 07-261-I, 1:2000) and mGlu3 (Alomone Labs AGC-012, 1:2000) were used as previously characterized (Sanger et al 2013; Yang et al 2017). After incubation with HRP-conjugated secondary antibody (Jackson Immuno; 1:20,000-1:40,000),immunoreactive bands on the membranes were detected by enhanced chemiluminescence (ECL Plus; GE Healthcare Bio-Sciences). For total protein expression, blots were re-probed with calnexin (Millipore AB2301 1:20,000-1:40,000) as a loading control (Johnson 2012). Band density was measured using NIH ImageJ software.

### Statistical analyses

All statistics were completed on SPSS (IBM, Armonk, NY) or Graphpad Prism (version 5.00, GraphPad Software, La Jolla, CA). Sidak’s post-hoc comparisons were used when required. An alpha level of p < 0.05 was set for all statistical analyses. In Experiment 1, band density of the protein of interest was averaged for the cocaine-naïve control group. The band density of individual cocaine rats was then divided by this control average to yield values that were expressed as % control (which was set to 100%). For the total protein fraction, this occurred after first adding together monomer and dimer expression of mGlu2 or 3 and then dividing this sum by the density of the calnexin band to control for the amount of protein loaded. For male rats, these values were compared with one-way analysis of variance (ANOVA). For female rats, a 2-way Phase (Estrus/Non-estrus) × Treatment (Cef/Veh) ANOVA was first conducted on the normalized data. Upon finding no effect of estrous cycle for any protein, the cycle data were collapsed, and a one-way ANOVA was conducted so that expression could be compared between the cocaine groups and the cocaine-naïve control group akin to the analysis in male rats. Monomer and dimer values were summed only for total protein expression, as only the dimer band was present in blots from the biotinylated fraction, consistent with the idea that only dimerized proteins form the active receptor present in the membrane. Tests for main effects of Group and Time, and Group × Time interactions were conducted using repeated measures ANOVAs on active and inactive lever presses for the self-administration and extinction phases of the experiments separately. For Experiment 2, behavioral data (self-administration, extinction and reinstatement test) were analyzed using multi-factorial Treatment (Cef/Veh) × NA core infusion (LY341495/Veh) × Time ANOVAs, with Time as a repeated measure (RM).

### Experiment 1 – The effects of cocaine and ceftriaxone on surface and total expression of mGlu2 and mGlu3 in the nucleus accumbens

We began this project by conducting western blots in female NA core tissue that remained from a previous project (Bechard et al 2018). We hypothesized that cocaine exposure decreases expression of mGlu2 and mGlu3 and that ceftriaxone restores this expression, as is seen with other proteins in the NA core. Therefore, we compared cocaine-naïve animals protein expression levels to animals that had self-administered cocaine and treated with vehicle and those treated with ceftriaxone. Ceftriaxone does not alter GLT-1 expression in the absence of exposure to cocaine (Knackstedt et al., 2010) Therefore, in this project, female rats underwent cocaine self-administration, extinction and Cef/Veh treatment as described above. These rats were killed, NA core tissue dissected and processed via the biotinylation protocol as described above. We used total protein and biotinylated fraction tissue from 16 vehicle-treated rats (8 in estrus; 8 non-estrus) and 15 ceftriaxone-treated rats (8 estrus; 7 non-estrus) and 14 cocaine-naïve control rats (7 estrus; 7 non-estrus). All of these rats were used in Bechard et al., 2018; no samples were excluded. We also included 15 male rats that underwent cocaine self-administration and extinction as described above; 8 received vehicle and 7 ceftriaxone. We also utilized 7 cocaine-naïve male rats. Tissue from the total and surface fraction were blotted for mGlu2 and mGlu3 as described above.

### Experiment 2 – The effect of intra-accumbens mGlu2/3 antagonist LY-341495 on the ability of ceftriaxone to attenuate cue- and cocaine-primed reinstatement

This experiment used 40 rats (27 females, 13 males). As shown in the experimental timeline (Fig. 4a), all rats underwent cocaine self-administration and 2 weeks of extinction training. Rats were treated with ceftriaxone (n=23) or vehicle (17) during the final 6 days of extinction training. Rats were not tested for reinstatement of cocaine-seeking during the estrus phase of the estrous cycle as ceftriaxone is not effective at attenuating reinstatement of cocaine-seeking during this phase, while it is effective in other stages (Bechard et al., 2018). Immediately prior to cue-primed reinstatement test, rats received bilateral intra-NA core infusion of the mGlu2/3 receptor antagonist LY341495 (n=18) or saline (n=22) via a Harvard Apparatus Pump at a flow rate of 0.25μL/minute. During the 2 hr cue-primed reinstatement test, lever presses on the previously active lever resulted in presentation of the cues previously paired with cocaine infusions, although no cocaine was available during this test. After the cue-primed reinstatement test, 24 females and 13 males underwent 2-3 additional days of extinction training until meeting extinction criteria. Three female rats (2 Veh; 1 Cef) underwent a cue-prime reinstatement test but not cocaine-primed test due to illness or failure to extinguish lever pressing following the cue-primed reinstatement test. Rats continued to receive ceftriaxone or vehicle during these 2-3 days. Prior to the 2 hr cocaine-primed reinstatement test, rats received bilateral infusions of either LY341495 (n=15) or saline (n=22) in a counterbalanced manner such that rats receiving vehicle during the cue test received LY during the cocaine-primed test. After LY administration, rats were injected with cocaine (10 mg/kg, IP) and placed into the operant chamber. During this test, lever presses were recorded but did not produce drug or cues. Following the cocaine-primed reinstatement test, animals were deeply anesthetized with pentobarbital (100 mg/kg, i.p.). Rats were transcardially perfused with phosphate-buffered saline (PBS) and then 4% paraformaldehyde (PFA). Brains were extracted and preserved in 4% PFA for 24 hr followed by 20% sucrose in PBS for 48 h. Brains were frozen and stored at −80° C until they were sliced at 20 μm using a cryostat to verify cannula placement.

## Results

### Experiment 1 – Cocaine decreases, and ceftriaxone increases mGlu2 expression but not mGlu3 expression

Self-administration data from the female rats used for western blotting were presented previously (Bechard et al., 2018). For male rats, self-administration behavior did not differ between rats later assigned to Cef/Veh (Fig. 1). There was no main effect of Treatment or Treatment × Time interaction on active lever presses during self-administration or during extinction (analyses conducted separately) in rats that later received Cef or Veh (Fig. 1a). There was an effect of Time on active lever presses during self-administration as both groups increased lever presses [F(11,45)=9.09, p<0.001]. There was a main effect of Time on active lever presses during extinction [F(11,484)=1.88, p=0.04] as both groups decreased lever presses. There was no effect of Treatment or Treatment × Time interaction on inactive lever presses during self-administration or extinction (analyses conducted separately; Fig. 1b). There was a main effect of Time on inactive lever presses during self-administration, as both groups decreased pressing after the first day of self-administration [F(11,506)=6.4, p<0.001]. Male rats later assigned to receive ceftriaxone or vehicle did not differ in cocaine intake (mg/kg) during cocaine self-administration (Fig. 1c). A significant main effect of Time was detected for cocaine intake [F(11,484)= 4.58, p<0.0001], as both groups increased intake over the 12 days.

**Fig. 1.**
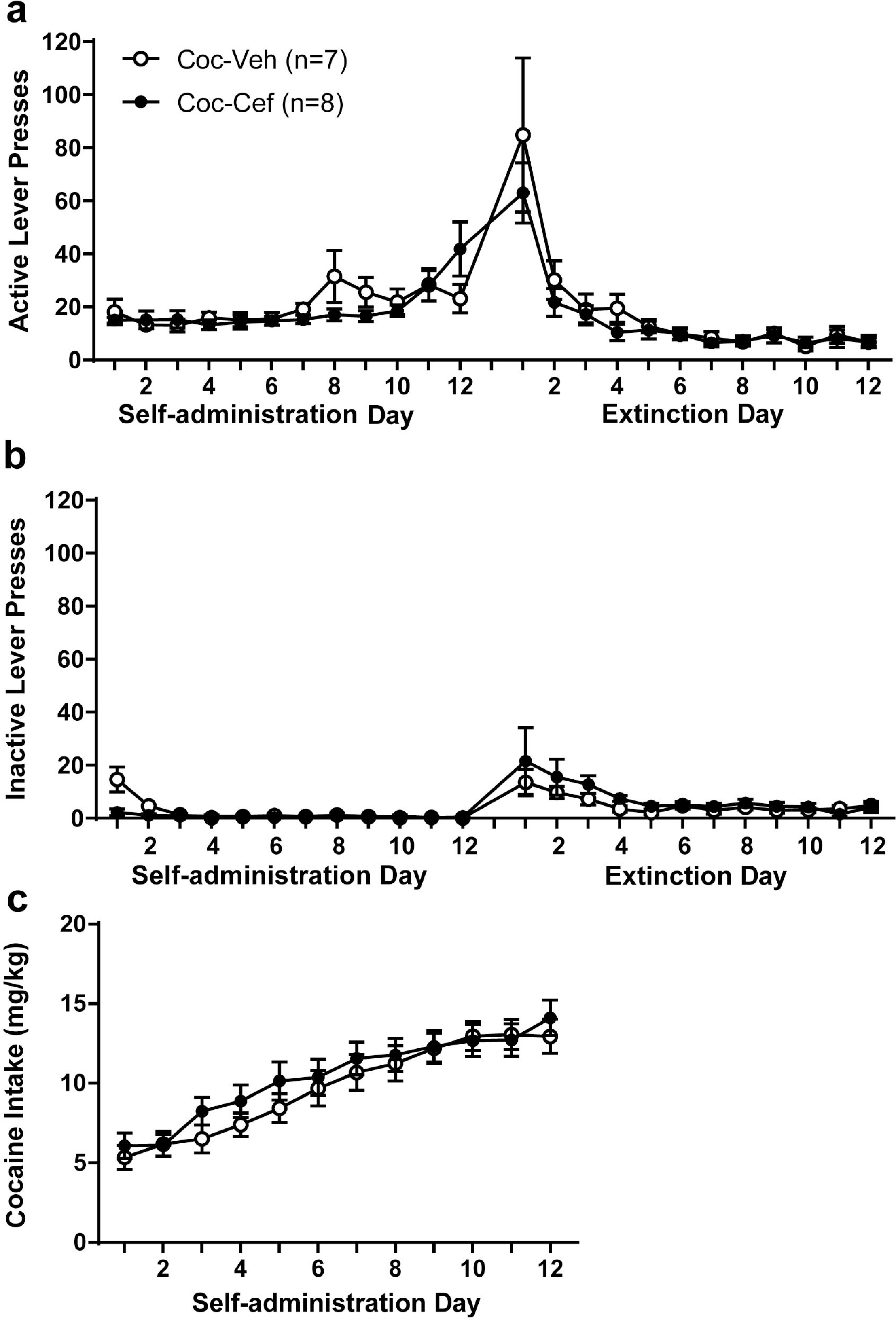
Self-administration behavior of rats utilized for western blotting did not differ between male rats later assigned to receive vehicle or ceftriaxone. Active **(a)** and inactive **(b)** lever presses during cocaine self-administration and extinction training, and **(c)** cocaine intake during self-administration did not differ between groups later treated with ceftriaxone/vehicle.

Total protein expression of mGlu2 differed by group for male rats [F(2,17)= 4.626, p<0.05], with post-hoc tests showing that the Coc-Veh group displayed reduced expression relative to both Control and Coc-Cef groups (Fig. 2a). Surface expression of mGlu2 also differed by group for male rats [F(2,19)= 5.334, p<0.05], however the Coc-Veh group only displayed reduced expression relative to the Coc-Cef group (Fig. 2c).

**Fig. 2.**
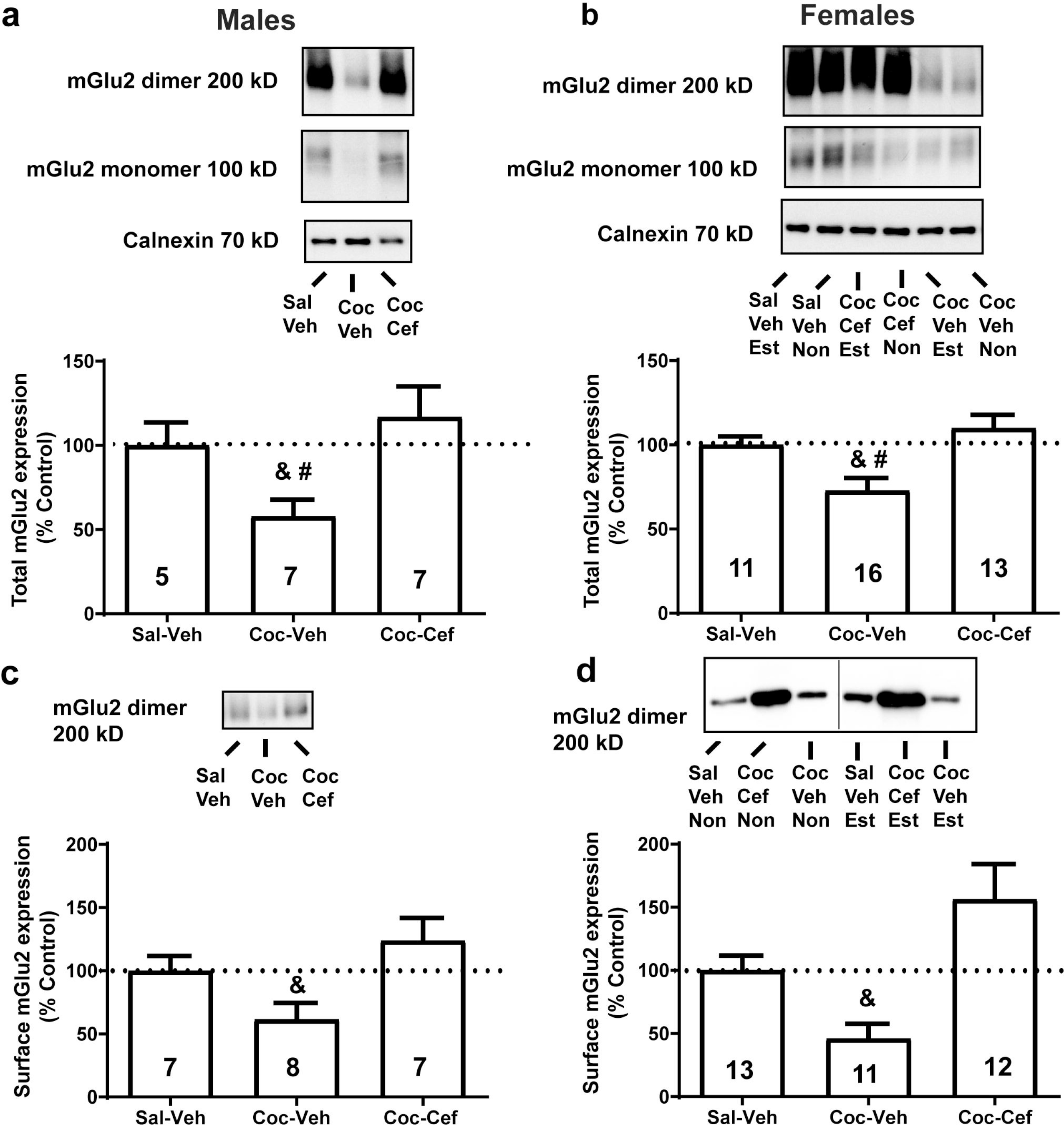
Cocaine decreases, and ceftriaxone increases surface and total mGlu2 expression in the NA core. **a** Cocaine decreases and ceftriaxone increases total protein expression of mGlu2 in the NA core of male rats. **b** Total protein expression of mGlu2 in the NA core is decreased by cocaine and restored by ceftriaxone in female rats. **c** Surface expression of mGlu2 in male rats is increased by ceftriaxone. **d** Surface expression of mGlu2 is increased by ceftriaxone in female rats. & = p<0.05 vs. Coc-Cef; # = p<0.05 vs. Sal-Veh.

When estrous cycle was considered, a two-way ANOVA conducted on total mGlu2 expression in female rats found a main effect of Treatment [F(1,25)= 3.918, p=0.05] but not estrous cycle Phase and no Treatment × Phase interaction. Since expression did not differ by estrous phase, we then collapsed across phase such that a one-way ANOVA could be conducted that included the control group. This analysis yielded a significant main effect for total mGlu2 in females [F(2,37)= 9.00, p<0.001]. Post-hoc tests found that as in males, the Coc-Veh group displayed reduced expression relative to both Control and Coc-Cef females (Fig. 2b). When a 2-way Treatment × Phase ANOVA was conducted, a significant main effect of Treatment was found for surface mGlu2 expression [F(1,19)= 8.875, p<0.01]. There was no main effect of estrous cycle Phase and no Treatment x Phase interaction. Thus, a one-way ANOVA was then conducted on surface expression of mGlu2, finding that as in male rats, surface mGlu2 also differed by group for female rats [F(2,33)= 8.826, p<0.001], with the Coc-Veh group only displaying reduced expression relative to the Coc-Cef group (Fig. 2d). No group differences in total or surface expression of mGlu3 were observed for male rats (Fig. 3a,c). No group differences in total or surface expression of mGlu3 were observed for female rats (Fig 3b,d). Sample size differed between analyses due to unreadable signal of some immunoreactive bands (due to loss of sample from gel) which were excluded from analysis.

**Fig. 3.**
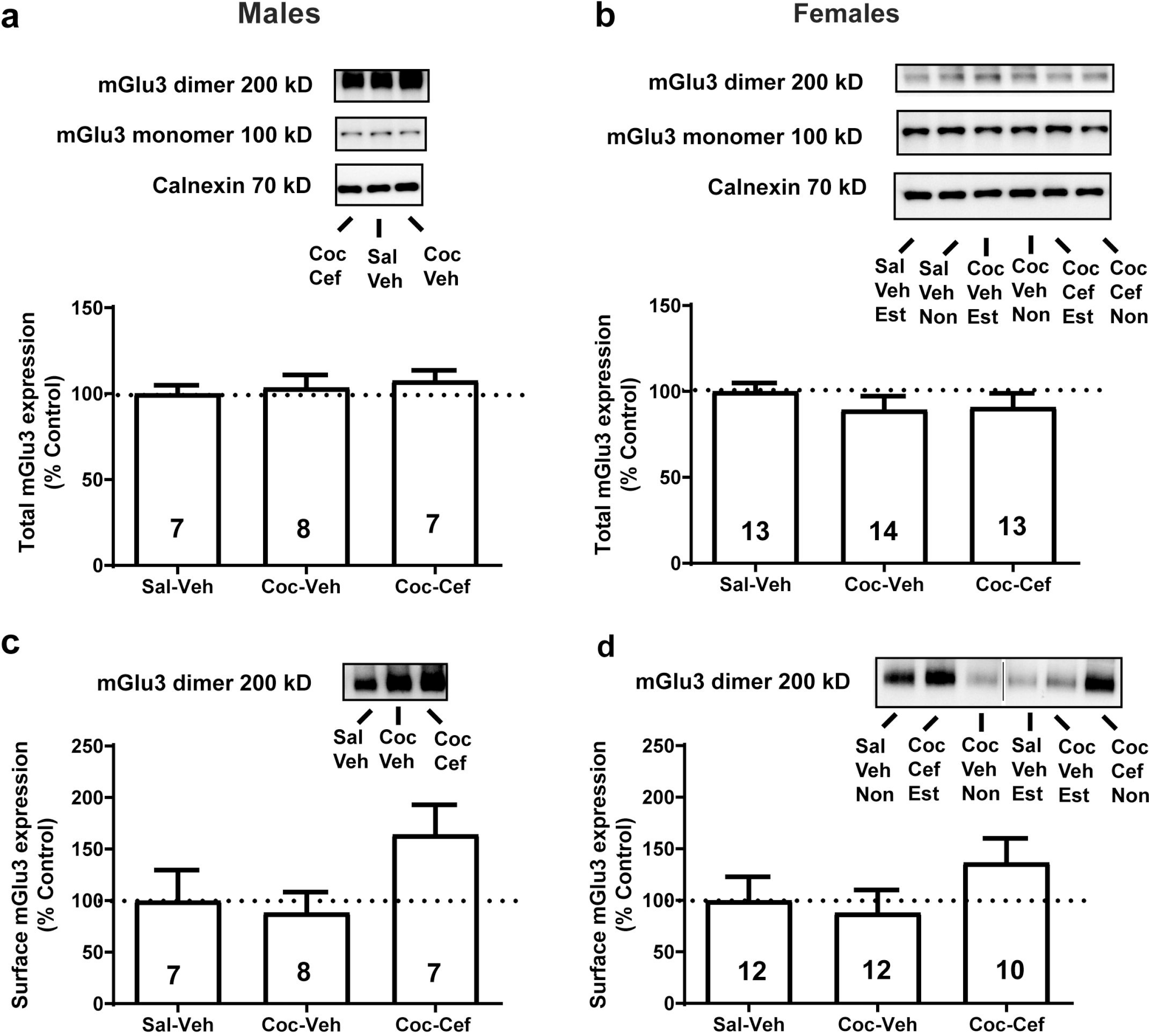
Cocaine and ceftriaxone have no effect on surface and total mGlu3 expression in the NA core. Total protein expression of mGlu3 was not altered by cocaine self-administration or ceftriaxone in male **(a)** or female **(b)** rats. Surface expression of mGlu3 was not altered by cocaine self-administration or ceftriaxone in male **(c)** or female rats**(d)**.

**Fig. 4.**
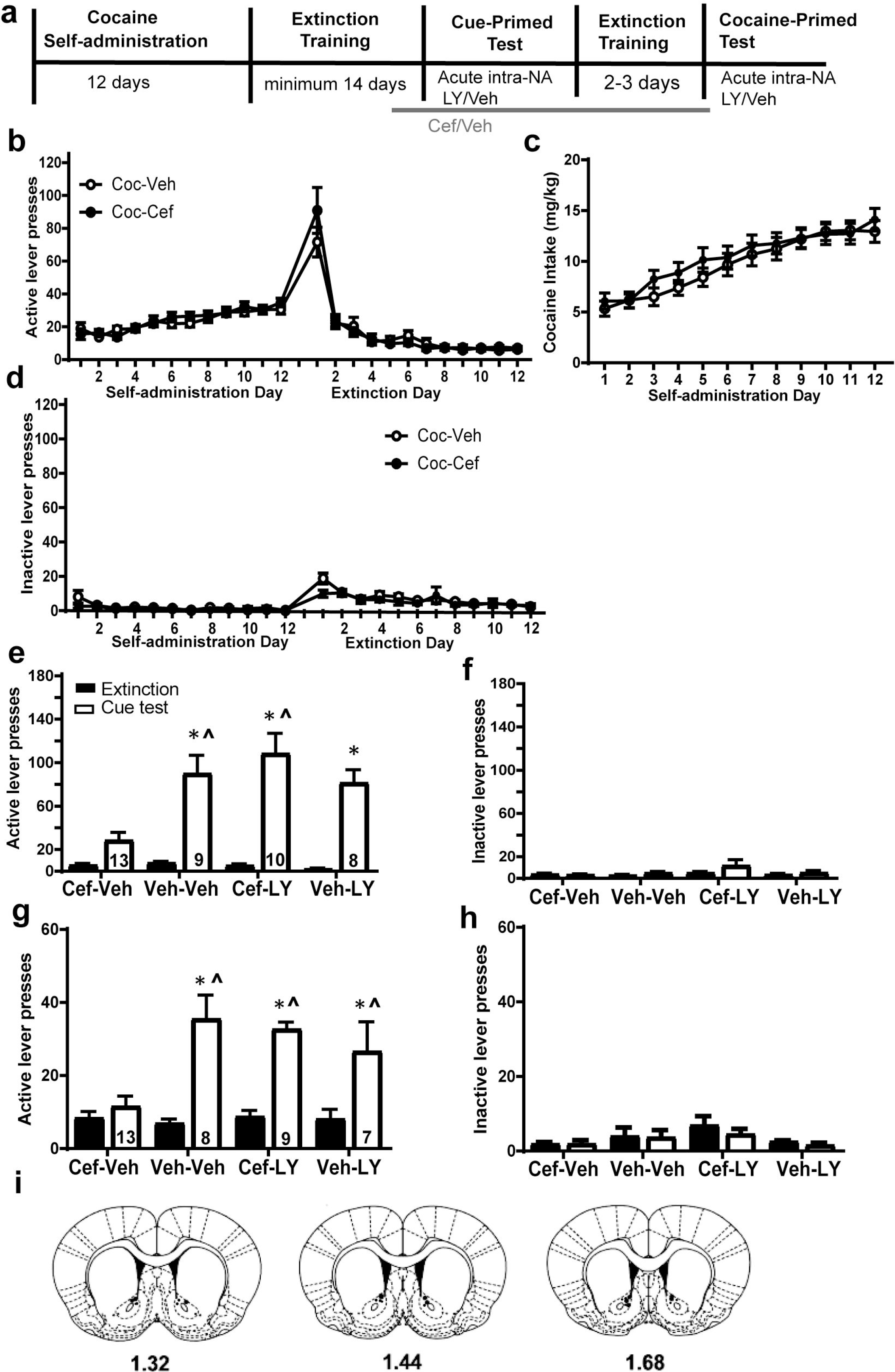
mGlu2/3 antagonism in the nucleus accumbens prevents ceftriaxone from attenuating cue- and cocaine-primed reinstatement of cocaine seeking. **a** timeline of Experiment 2. Active lever presses during self-administration **(b)** and cocaine intake (mg/kg) during self-administration **(c)** did not differ between treatment groups. **d** Inactive lever presses during cocaine self-administration and extinction training also did not differ between treatment groups. **e** Presses on the previously active lever during the final days of extinction training and during a cue-primed reinstatement test. Active lever pressing increased during the test in Veh-Veh, Veh-LY, and Vef-LY groups; the Cef-Veh group did not reinstate drug-seeking. **f** Inactive lever presses during the cue-primed reinstatement test did not increase for any treatment groups compared to extinction. **g** Presses on the previously active lever during extinction training and a cocaine-primed reinstatement test. Lever presses increased during the test in Veh-Veh, Veh-LY, and Cef-LY treatment groups only; the Cef-Veh group did not reinstate drug-seeking. **h** There was no change in inactive lever presses between extinction and the cocaine-primed test. **i** Location of intracranial injector tips in the NA core as determined using the rat brain atlas (Paxinos and Watson, 2007). * = p<0.05 vs. Extinction. ^ = p<0.05 relative to Cef-Veh.

### Experiment 2: Intra-NA core infusion of LY341495 prevents ceftriaxone from attenuating reinstatement of cocaine-seeking

40 rats (males n= 13, females n= 27) were implanted with bilateral cannula and jugular catheters during surgery. Rats underwent behavioral testing according to the timeline in Fig. 4a. No main effect of Group and no Group × Time interaction were detected for active lever presses during self-administration and extinction (analyses conducted separately) in rats that later received Cef or Veh (Fig. 4b). There was an effect of Time [F(11, 220)= 10.03, p<0.0001] on active lever presses during self-administration as all groups increased lever pressing. There was an effect of Time on active lever presses during extinction training [F(11,407)= 50.88, p<0.0001], as all groups decreased presses on the previously active lever. No main effect of Group or Group × Time interaction was found for cocaine intake during self-administration for rats later treated with Cef or Veh (Fig. 4c). There was an effect of Time on cocaine intake [F(11,407)= 39.64, p<0.0001], as all groups increased intake throughout self-administration. Male and female rats did not differ in cocaine intake (mg/kg) during self-administration (not shown). There was no effect of Group or Group × Time interaction on inactive lever presses during self-administration or extinction training (analyses conducted separately; Fig. 4d). There was an effect of Time [F(11, 396)= 2.986, p=0.0008] on inactive lever presses during self-administration as all groups exhibited increased lever presses on the inactive lever presses early in self-administration. There was a main effect of Time on inactive lever presses during the first 12 days of extinction [F(11,396)=7.601, p<0.0001] as all groups pressed the inactive lever more in early extinction training.

Reinstatement of drug-seeking is defined by an increase in presses on the previously active lever during the reinstatement test compared to the average presses during the final two days of extinction training (Fig. 4e). A three-way Treatment × NA core infusion × Time interaction was detected for presses on the previously active lever during extinction and test [F(1,35)= 4.83, p=0.035]. There were no differences in lever pressing during the final days of extinction training between the four groups (p>0.05). Rats that were treated with Cef and an intra-NA core infusion of vehicle (e.g. Cef-Veh) did not increase lever pressing during a cue-primed reinstatement test compared to lever presses during the final days of extinction training (p=0.4). All other treatment groups increased lever pressing during a cue-primed reinstatement test compared to lever presses during the final days of extinction training systemic vehicle, intra-accumbal vehicle (Veh-Veh; p<0.001), systemic vehicle, intra-accumbal LY341495 (Veh-LY; p=0.04), systemic Cef, intra-NA core LY341495 (Cef-LY; p<0.001). Lever presses during a cue-primed reinstatement test did not differ between rats treated with Veh-Veh and those treated with Veh-LY or Cef-LY, but Cef-Veh rats displayed reduced lever pressing relative to both Veh-Veh and Cef-LY groups. Thus, an intra-NA core infusion of LY341495 immediately prior to a cue-primed reinstatement test prevented ceftriaxone from attenuating reinstatement of cocaine-seeking (Fig. 4e). For inactive lever presses during extinction and cue-primed reinstatement test, a 3-way interaction was not detected, nor were any 2-way interactions detected (Fig. 4f)

While a 3-way interaction was not detected for active lever pressing during extinction and cocaine primed reinstatement, a Treatment × Time interaction was detected [F(3,29)=5.84, p= 0.003]. There was also a Treatment × NA core infusion interaction [F(1,35)= 3.36, p=0.002], indicating that LY infusion has different effects in Cef and Veh-treated rats. Rats treated with Veh-Veh (p<0.001), Veh-LY (p=0.01), and Cef-LY (p<0.001) displayed a significant increase in lever pressing from extinction to test. Rats that were treated with Cef-Veh did not increase lever presses on the previously active lever during the cocaine-primed reinstatement test. Similar to the cue-primed reinstatement test, Cef-LY rats did not differ from rats treated with Veh-Veh in lever presses during a cocaine-primed reinstatement test, but Cef-Veh rats exhibited less presses than all other groups during the test (p’s<0.05). Thus, intra-NA core LY341495 immediately prior to the cocaine-primed reinstatement test prevented ceftriaxone from attenuating reinstatement of cocaine-seeking (Fig. 4g). There were no three-way or 2-way interactions detected for inactive lever presses during extinction and the cocaine-primed reinstatement test (Fig. 4h). See Fig. 4i for placement of injector tips in the NA core for LY341495 or vehicle infusion. While all injector tips were located in the NA core, the proximity to the lateral ventricles cannot rule out the possibility that spread to the ventricles occurred. However, the small infusion volume (0.5 μL) likely prevented that possibility.

## Discussion

Here we present data that for the first time implicates NA core mGlu2/3 expression and function in the ability of ceftriaxone to attenuate the reinstatement of cocaine seeking in male and female rats. While neither cocaine self-administration nor ceftriaxone produced changes in mGlu3 total or surface expression in the NA core, mGlu2 total and surface expression was decreased by cocaine self-administration and increased by ceftriaxone in both male and female rats. This upregulation of mGlu2 receptors may be a key aspect of how ceftriaxone attenuates reinstatement of cocaine-seeking. Furthermore, we demonstrate that mGlu2/3 signaling in the NA core is necessary for ceftriaxone to attenuate cue-and cocaine-primed reinstatement.

Both mGlu2 and mGlu3 receptors are expressed abundantly in the NA core where mGlu3 receptors are found in both pre- and post-synaptic cells and also on glial processes (Tanabe et al 1993; Ohishi et al 1993; Petralia et al 1996; Tamaru et al 2001). mGlu2 receptors, however, are found specifically on presynaptic terminals (Ohishi et al. 1994). Here, we infused an mGlu2/3 antagonist into the NA core where it would bind with similar efficacy to both mGlu2 and mGlu3 receptors. While we speculate that the antagonism of presynaptic mGlu2 autoreceptors underlies the ability of LY341495 to block ceftriaxone’s effects on reinstatement, at this time, involvement of mGlu3 cannot be ruled out. The changes in mGlu2 total and surface expression and lack of change in mGlu3 in the current study further suggest that ceftriaxone interacts with mGlu2 to attenuate cocaine seeking. The same pattern of mGlu2 and 3 changes were observed in males and females, regardless of estrous cycle phase in females. Thus, the inability of ceftriaxone to attenuate reinstatement when females are in estrus (Bechard et al., 2018) is not due to different levels of total or surface mGlu2 expression in estrus. The ability of ceftriaxone to upregulate GLT-1 and xCT is also not prevented by estrus (Bechard et al 2018), raising the possibility that reinstated cocaine-seeking during estrus is mediated by another neurotransmitter in NA core (e.g. dopamine) or another brain region. Estrus-induced adaptations throughout the rodent brain require investigation to better understand how sex hormone fluctuations affect the reinstatement of cocaine-seeking.

It has been suggested that the loss of NA core mGlu2/3 binding and function is due to the reduced basal levels of glutamate present following cocaine self-administration (Baker et al., 2002, Moussawi et al., 2011). Here we detected a reduction in mGlu2 expression not just in the surface membrane fraction, but also in the total protein fraction. While our results do not rule out a role for basal glutamate in mediating mGlu2 adaptations after cocaine, changes in total protein expression may indicate that cocaine reduces mGlu2 expression through a transcriptional or trafficking mechanism such as a reduction in mGlu2 mRNA in regions projecting to the NA core. Previous work has identified distinct and overlapping patterns of expression for mGlu2 and mGlu3 mRNA expression. mGlu3 mRNA is found in the NA core (McOmish et al., 2016) while mGlu2 mRNA is not present in the NA core, consistent with this receptor’s presence only on terminals arising from afferent regions (Ohishi et al., 1993). mGlu2 mRNA is present in cerebral cortical brain regions and the amygdala, regions which send projections to the NA core (Ohishi et al., 1994; Testa et al., 1998). We speculate that cocaine decreases, and ceftriaxone increases, mGlu2 mRNA in the PFC and other brain regions that send glutamate projections to the NA core. This possibility will be examined in future studies. We did not assess the effects of ceftriaxone on mGlu2/3 expression in cocaine-naïve rats, as we find no effect on xCT and GLT-1 expression, nor any effect on basal glutamate, in the absence of cocaine intake (Knackstedt et al 2010). Thus, the upregulation of mGlu2 and 3 expression by ceftriaxone in the absence of cocaine would implicate a transcriptional/translational mechanism in this effect. Evidence suggests that ceftriaxone works through the nuclear factor-kappaB (NF-kappaB) signaling pathway (Lee et al 2008). This pathway has also been implicated in transcriptional regulation of mGlu2 (Chiechio et al 2006), providing a potential mechanism through which ceftriaxone increases mGlu2 expression.

While the current work suggests that upregulation of mGlu2 receptors in the NA core is necessary for ceftriaxone to attenuate reinstatement of cocaine-seeking due to their ability to regulate glutamate efflux during reinstatement, ceftriaxone’s effects on dopamine signaling in the NA core has also not yet been examined. The system xc- antagonist CPG increases dopamine levels in the NA core, an effect that is blocked by an mGlu2/3 agonist (Baker et al., 2002(Xi et al 2002). The mGlu2/3 agonist L-CCG-1 reduces KCl-evoked dopamine release in the NA core (Chaki et al., 2006). Thus, the antagonist LY341495 may have also increased NA core dopamine.

In conclusion, the present results indicate that mGlu2/3 signaling in the NA core is necessary for ceftriaxone to attenuate the reinstatement of cocaine seeking. mGlu2, and not mGlu3, total and surface expression was reduced by cocaine and restored by ceftriaxone. Because mGlu2 is found on presynaptic glutamate terminals in the NA core, it is likely that ceftriaxone works to upregulate these receptors and restores the ability to regulate presynaptic glutamate release after presentation of drug-associated cues and cocaine itself to reduce cocaine seeking. Future work will test the possibility that mGlu2 expression in projections from the PFC to the NA core is necessary for ceftriaxone to attenuate reinstatement and the glutamate efflux that accompanies it.

## Acknowledgements

The authors would like to thank Megan Milner, Matthew Molk and Tanuj Prajapati for outstanding technical assistance that facilitated the collection of data.

